# Tardigrades survive reversible dormancy during spaceflight: In-orbit revival, post-flight recovery, and transcriptomic responses

**DOI:** 10.64898/2026.11.06.764766

**Authors:** Divya Vishnu, Shravani Pillanjinayya, Harikumar R Suma, Swati Ghosh, Jayakumar Pallavi, Krishnan Yuvashree, Vrishali Murugkar, Hitesh Tikariha, Gautam Das, Vibhor Singh, Sandeep M Eswarappa

## Abstract

Long-duration space missions face substantial challenges, including limited resources and prolonged exposure to microgravity and radiation. Transporting organisms in a dormant state and reviving them at their destination could help overcome some of these limitations. However, the feasibility of this strategy has not been experimentally demonstrated in multicellular animals beyond Earth. Here, we investigated this possibility using tardigrades, microscopic invertebrates capable of entering a reversible state of dormancy. We demonstrate that dormant tardigrades can withstand near-absolute-zero temperatures (∼12 mK), near-vacuum conditions (10^-9^ mbar), high doses of ionizing radiation (3000 Gy), and complete desiccation. We transported dormant tardigrades to the International Space Station (ISS). They were revived aboard the ISS, subsequently returned to a dormant state, and successfully revived again following their return to Earth. Approximately 45% of the space-flown tardigrades survived the 64-day experimental period. This included two dormancy-revival cycles, 18 days aboard the ISS, and prolonged periods without food or water. Importantly, surviving tardigrades retained their reproductive capacity. Transcriptomic analysis of ISS-exposed tardigrades revealed upregulation of genes associated with protein folding and stress responses, accompanied by downregulation of genes involved in energy metabolism and protein synthesis, providing insights into their molecular responses to the spaceflight environment. Together, these findings demonstrate the successful revival and re-induction of dormancy in a multicellular animal beyond Earth and establish an experimental foundation for exploring dormancy-based strategies for future long-duration space missions.

## Introduction

Travel from Earth to Mars using current propulsion technologies requires approximately seven to ten months [1], and missions to more distant destinations would involve even longer durations. Several ongoing space programs aim to send human astronauts to the Moon in the near future and to Mars in the following decades (e.g., NASA’s Artemis program). Such long-duration missions present major logistical challenges, including sustained provision of food, water, and oxygen. They also expose organisms to prolonged microgravity and ionizing radiation, which are principal biological constraints of deep-space travel. In addition, if exposed directly to the external space environment, organisms will encounter near-vacuum conditions, extreme thermal fluctuations, and ultraviolet radiation, all of which are lethal [2]. These challenges make long-duration interplanetary transport of astronauts or even model organisms challenging with current technologies.

One potential solution to the logistical challenges of long-duration space travel is to transport model organisms (or even astronauts) in a dormant (or hibernation-like) state and revive them upon arrival at their destination. Owing to their greatly reduced metabolic activity, dormant or hibernating organisms require substantially fewer resources, including food, water, and oxygen. Consequently, considerable efforts are underway to develop approaches for inducing such states in mammals [3–5]. However, despite its conceptual appeal, the revival of dormant animals after transport beyond Earth has not yet been experimentally demonstrated.

Tardigrades are microscopic invertebrates that provide a compelling model to explore this strategy. Limno-terrestrial tardigrades can enter a reversible, ametabolic dormant state known as the “tun” state following slow and controlled dehydration [6, 7]. They remain viable for months to years in this state until rehydration. In this state, tardigrades tolerate otherwise lethal conditions, including extreme temperatures and pressures, ionizing and ultraviolet radiation, and certain conditions encountered in low Earth orbit (TARDIS and LIFE-TARSE experiments on FOTON-M3) [8–17].

In this study, we investigated whether the eutardigrade *Paramacrobiotus* sp. could be transported to the ISS in a dormant state, revived in orbit, re-induced into dormancy, and subsequently revived upon return to Earth. We assessed the long-term survival and reproductive capacity of tardigrades following ISS exposure. We also studied their transcriptomic responses to ISS environment through comparative RNA sequencing of ISS-exposed and ground-control animals.

## RESULTS

### Dormant tardigrades withstand multiple extreme physical conditions found in space

In our previous studies, we reported that a newly identified tardigrade, *Paramacrobiotus* sp. BLR strain could survive UV-C radiation at doses up to 4000 J/m² and temperatures up to 85°C [17, 18]. Morphological characterization and sequencing of the mitochondrial *COI* (cytochrome c oxidase subunit I) and nuclear ITS2 (internal transcribed spacer 2) regions indicated that the organism belongs to the genus *Paramacrobiotus* [18]. To determine its precise taxonomic identity, we generated a genome assembly using a hybrid approach combining Nanopore and Illumina reads (GenBank accession # GCA_052674745.1) (Table S1 and S2). Identity analysis of conserved marker genes using BLASTn revealed that they are less than 90% identical to their best match (except rRNAs), which were from *Paramacrobiotus metropolitanus* Sugiura, Matsumoto & Kunieda, 2022 [19] (GenBank accession # GCA_019649055.1) (Table S3). Phylogenetic analysis based on 74 BUSCO (Benchmarking Universal Single-Copy Orthologs) marker genes showed that the BLR strain forms a genomically distinct lineage within the genus *Paramacrobiotus.* Based on this phylogenomic distinctiveness, we designated this tardigrade as *Paramacrobiotus karunaadensis*, named after the southern region of India from where it was isolated (Fig. 1A,B).

**Figure 1.**
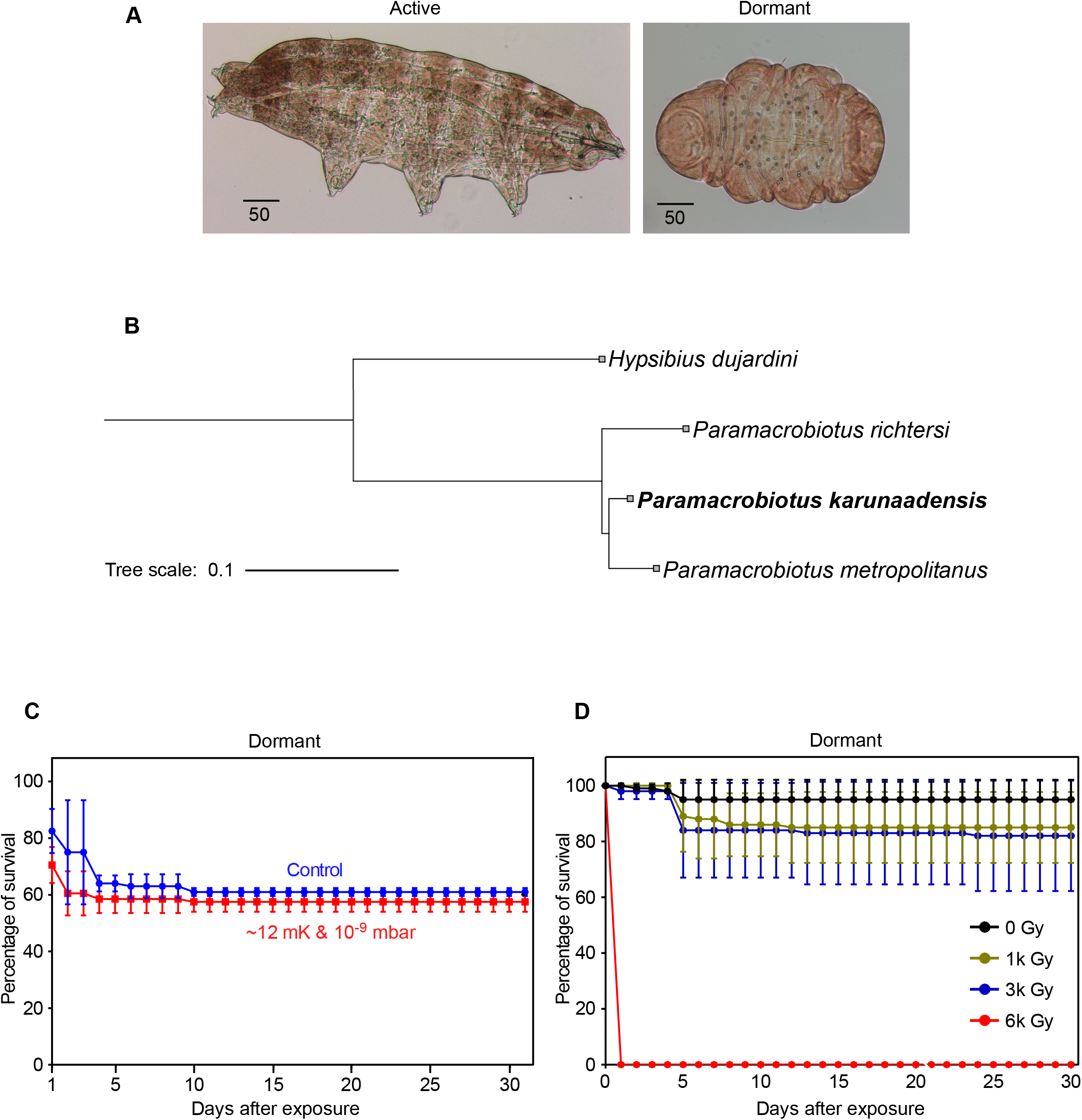
Tolerance of dormant tardigrades to extremely low temperatures, near-vacuum conditions, and ionizing radiation. (A) Microscopic images of *P. karunaadensis* tardigrades in active (left) and dormant state (right). Scale bar, 50 μm. (B) Phylogenomic tree of select tardigrade species based on 74 BUSCO marker genes. BUSCO phylogenomic tool was used to construct the tree. All genes considered here are complete and found in a single copy in all input samples. (C) Survival of dormant tardigrades exposed to 12 mK temperature and 10^-9^ mbar pressure inside a dilution refrigerator for 24 hours. N = 50 tardigrades in each group. The difference in survival is not statistically significant (P > 0.05, Log rank test). -▪-, tardigrades exposed to ∼12 mK & 10^-9^ mbar and -●-, tardigrades exposed to standard laboratory conditions (Control). (D) Survival of dormant tardigrades exposed to multiple doses of gamma radiation. N = 50 tardigrades in each group. 0,1, and 3 kGy groups are statistically significant from the 6 kGy group (P < 0.05, Log rank test). Graphs in (C) and (D) show data from two independent experiments performed using 50 tardigrades in each group. Error bars indicate SE.

Given its very high tolerance to UV radiation and desiccation, we examined whether *P. karunaadensis* could also survive and recover from exposure to some of the extreme environmental conditions relevant to astrobiology. Upon slow dehydration, tardigrades contract longitudinally and retract their head and limbs to form a reversible, ametabolic dormant structure known as the anhydrobiotic “tun” state, similar to other limno-terrestrial tardigrades (Fig. 1A) [15, 16]. In this state, tardigrades are known to tolerate a wide range of physical extremes. When gradually rehydrated, they return to their active form.

We exposed *P. karunaadensis* individuals in the tun state to an ultralow temperature of ∼12 mK (−273.14°C) and ultra-low pressure of 10^-9^ mbar (10⁻^12^ bar) inside a dilution refrigerator for 24 hours. Survival of dormant tardigrades under these conditions did not differ notably from that of controls maintained under standard laboratory conditions. After 30 days, 58 ± 4% of the exposed tuns were alive, compared with 61 ± 1% in the control group maintained in laboratory conditions (20°C and 0.91 bar) (Fig. 1C). In contrast, none of the active (non-dormant) tardigrades survived exposure to these extreme conditions. We next assessed tolerance to ionizing radiation. Approximately ∼82% of the dormant *P. karunaadensis* individuals survived exposure to 3000 Gy of γ-radiation (Fig. 1D), demonstrating resistance to multiple extreme stressors. Notably, the experimental conditions tested here are orders of magnitude more extreme than those typically encountered in space, where the temperature is approximately 2.7 K, and the radiation dose is in the milli-gray range [20, 21]. Together, the findings described above demonstrate that *P. karunaadensis* can survive a wide range of extreme physical stresses, including extreme cold (almost zero K), near-vacuum pressure, complete desiccation, lethal doses of ionizing radiation, and germicidal UV-C radiation (our previous study). These observations broadly define the tolerance limits of *P. karunaadensis*.

### Dormant tardigrades can be activated and re-induced back to dormancy at ISS

Based on these observations, *P. karunaadensis* was selected for the International Space Station (ISS) experiment conducted as part of the ISRO-NASA (Indian Space Research Organization-National Aeronautics and Space Administration) joint mission (Axiom-4) to the ISS, which operates in low Earth orbit (Fig. 2A). The primary objective of this mission was to test whether dormant tardigrades could withstand the cumulative stresses associated with transport to and from the ISS (launch, transport, residence aboard the ISS), and whether they could be successfully reactivated beyond Earth at ISS, returned to dormancy under ISS conditions, and subsequently revived after return to Earth.

**Figure 2.**
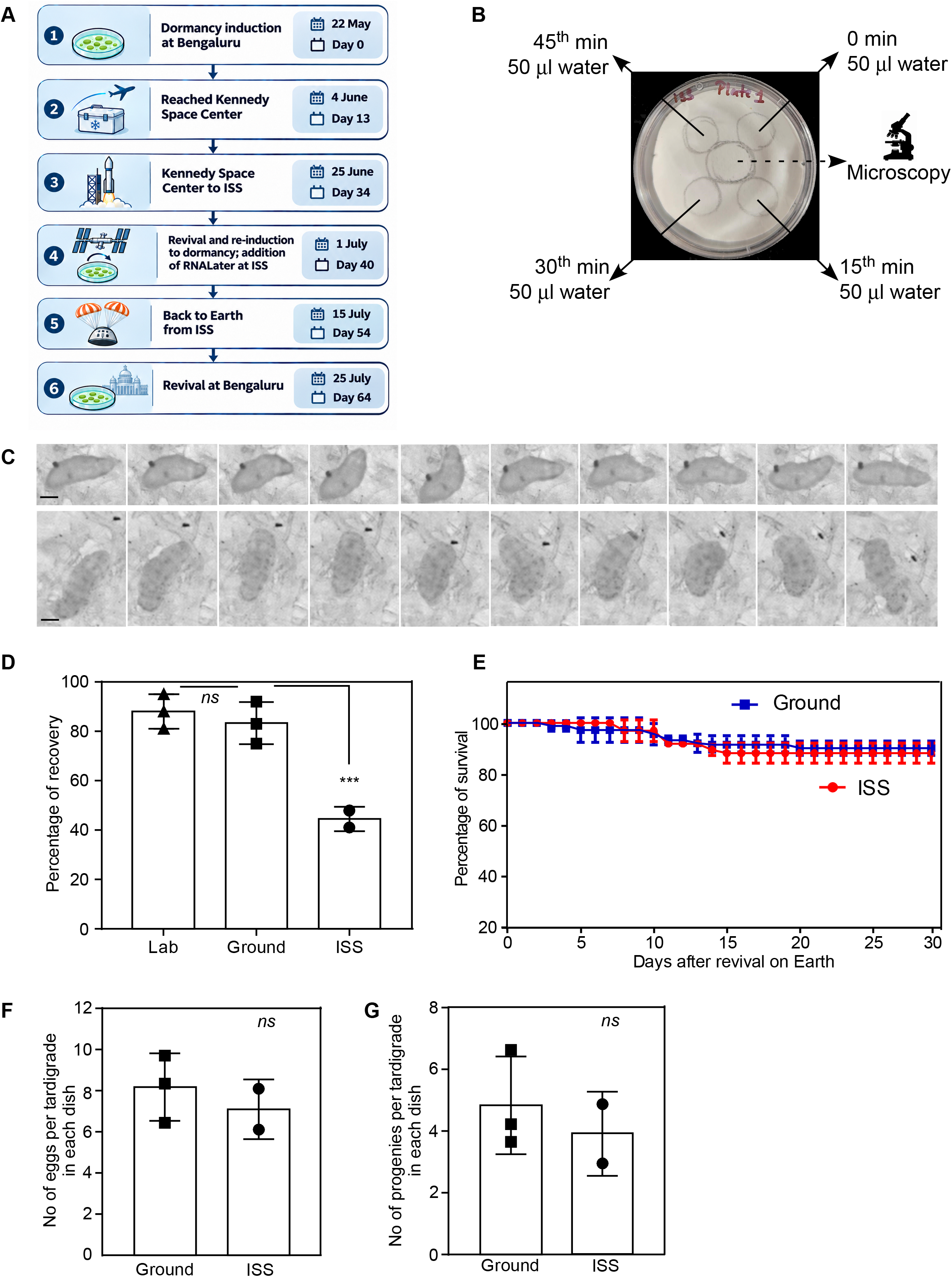
Reversible dormancy of tardigrades during spaceflight: In-orbit revival and post-flight recovery. (A) Timeline of the tardigrade spaceflight experiment, including preflight preparations, onboard ISS procedures, and post-flight recovery. All dates refer to 2025. (B) A representative image of the 35 mm dish used to carry dormant tardigrades to ISS. The locations where water was added for reactivation are indicated by circles. The central circle represents the place where dormant tardigrades were placed. (C) Sequential screengrabs (20 s intervals) showing the activation of tardigrades from dormancy aboard ISS. Images of two representative tardigrades are shown. Scale bar, 100 μm. More moving tardigrades can be seen in Video S1. (D) Percentage of dormant tardigrades reactivated on Earth. Dormancy was induced at ISS. Lab control results were obtained from the science verification test (SVT). ***, *P* = 0.0076. ns, not significant (*P* > 0.05). (E) The survival of these tardigrades for 30 days after reactivation on Earth (F) The egg-laying ability (expressed as number of eggs per recovered tardigrade in each dish) of these reactivated tardigrades was assessed over a two-month period. ns, not significant (*P* = 0.51). (G) Number of offspring produced by reactivated tardigrades (expressed as number of offspring per recovered tardigrade in each dish) over a two-month period. ns, not significant (*P* = 0.55). In all graphs, tardigrades from 2 (ISS) or 3 (Ground and Lab) Petri-dishes (Each had ∼90 individuals), their eggs, or their offspring are shown. Each point represents a Petri dish. -▪-, Ground control and -●-, ISS samples. Error bars indicate SE. *P* values were calculated using unpaired two-tailed t-tests with Welch’s correction.

Dormancy (anhydrobiotic “tun” state) was induced on day 0 by controlled dehydration, as described previously [18], on circular Whatman No. 3 filter paper discs (33 mm in diameter) in our laboratory (Methods). The filter papers containing dehydrated tardigrades were placed inside six 35 mm Petri dishes (∼90 individuals per disc) and shipped to the Kennedy Space Center, USA, where they were stored at 20°C until launch. Three dishes were designated for the ISS experiment, and three were ground controls. The ground-control samples were transported to the Kennedy Space Center like the flight samples, thereby undergoing identical storage duration and transport logistics, except for launch and orbital exposure. Experimental steps such as rehydration and re-induction of dormancy were performed simultaneously for both ISS and ground-control samples, with timing coordinated in real time during astronaut operations aboard the ISS.

The ISS samples were launched on day 34 (relative to the day of dormancy induction) aboard a SpaceX Falcon 9 Block 5 rocket carrying the Axiom-4 crew and experimental payload to low Earth orbit. The spacecraft subsequently docked with the ISS, where the samples remained for the duration of the 18-day mission.

On day 40 (sixth day aboard the ISS), the ISRO astronaut (Capt. Shubhanshu Shukla) activated the dormant tardigrades by gently adding 50 μL of sterile water around the filter paper region containing the specimens four times as described in Methods (Fig. 2B). About 1 hour after the final addition, the samples were observed under the onboard Keyence BZX800 microscope. The videos (Brightfield monochrome, 40 frames per second) of the activated tardigrades were recorded from five fields containing actively moving tardigrades (Fig. 2C, Video S1). In total, 106 (∼59%) of 180 tardigrades observed in the video exhibited clear movement within 15 min of rehydration. Because tardigrades can take several hours to resume full motility after rehydration, the actual recovery rate is likely higher than observed. These results demonstrate that *P. karunaadensis* can successfully be reactivated aboard the ISS. Nonetheless, we acknowledge that, because of the limited time available in the constrained environment of ISS, this observation is a qualitative demonstration of rehydration-induced activation rather than a quantitative functional assessment. Quantitative evaluation was therefore performed after return to Earth, as described below.

Following reactivation and imaging, the tardigrades were allowed to dehydrate under ISS environmental conditions, returning to the dormant tun state. The specimens spent a total of 18 days aboard the ISS, during which they were exposed to microgravity and elevated levels of cosmic and solar radiation relative to Earth.

### Dormant tardigrades exhibit partial recovery after transport to and from the ISS

The samples were returned to Earth on day 54 (relative to first dormancy induction). Dormant tardigrades recovered from the ISS were activated in our laboratory on day 64 using the same controlled rehydration protocol. We recovered a total of 78 active tardigrades, corresponding to ∼45% recovery, whereas the ground-control group (exposed to all conditions as ISS samples except spaceflight and ISS-associated microgravity) showed 83% recovery under identical conditions. The laboratory control group (maintained in our laboratory under optimal conditions) showed 88% recovery (Fig. 2D).

Reactivated tardigrades were subsequently monitored to determine whether space travel produced any long-term effects on survival or reproductive performance. Long-term survival was calculated for each group as the percentage of reactivated individuals that remained alive at the designated observation time relative to the total number of individuals initially reactivated. The survival percentage of the ISS group did not differ significantly from that of the ground-control group (Fig. 2E).

Reproductive performance was evaluated using two endpoints: egg production and second-generation offspring production. Egg production was calculated for each dish as the total number of eggs laid divided by the number of tardigrades. There was no statistically significant difference between ISS and ground control groups (Fig. 2F). Similarly, second-generation offspring production was calculated as the total number of offspring generated divided by the number of tardigrades in the corresponding dish. Second-generation offspring were observed in both the ISS and ground-control groups, with no statistically significant difference in the number of offspring produced per founder between the two groups under the present experimental conditions (Fig. 2G).

### Tardigrades exhibit transcriptional reprogramming aboard the ISS

As described above, the ISS environment exposes organisms to multiple stressors, including microgravity and elevated levels of ionizing radiation. To investigate the transcriptional responses of tardigrades to the ISS environment, we performed transcriptomic analysis of tardigrades exposed to spaceflight conditions. Our ISS payload included three tubes containing ‘active’ tardigrades, all from the same batch. Three additional similar tubes containing tardigrades were maintained as ground controls. Along with the survival and revival controls described above, these samples were transported to the Kennedy Space Center and returned. Consequently, they experienced transit and environmental conditions identical to those of the International Space Station (ISS) samples, excluding spaceflight-and ISS-specific factors.

On day 40 of the experiment (the sixth day aboard the ISS), ISRO astronaut Capt. Shubhanshu Shukla added RNAlater, an aqueous RNA-stabilizing reagent, to the ISS samples to preserve RNA integrity. RNAlater was simultaneously added to the ground-control samples in our laboratory. Following RNA stabilization, both the ISS and ground-control samples were stored at −80°C. These ISS samples were returned to our laboratory in frozen condition on day 54 for subsequent transcriptomic analysis.

Total RNA was isolated from the ISS and ground-control samples and subjected to next-generation RNA sequencing to investigate differential gene expression. De novo transcriptome assembly was performed using Trinity (v2.x) with paired-end sequencing reads. The assembled transcripts were functionally annotated using the Trinotate pipeline, incorporating BLAST-based homology searches against Swiss-Prot/UniProtKB and protein domain identification using HMMER. Principal component analysis (PCA) of the normalized transcript abundance data revealed distinct clustering of the three ISS samples, clearly separating them from the ground-control samples (Fig. 3A). This separation indicates substantial differences in gene expression profiles between the two groups, consistent with transcriptional alterations associated with exposure to the ISS environment.

**Figure 3.**
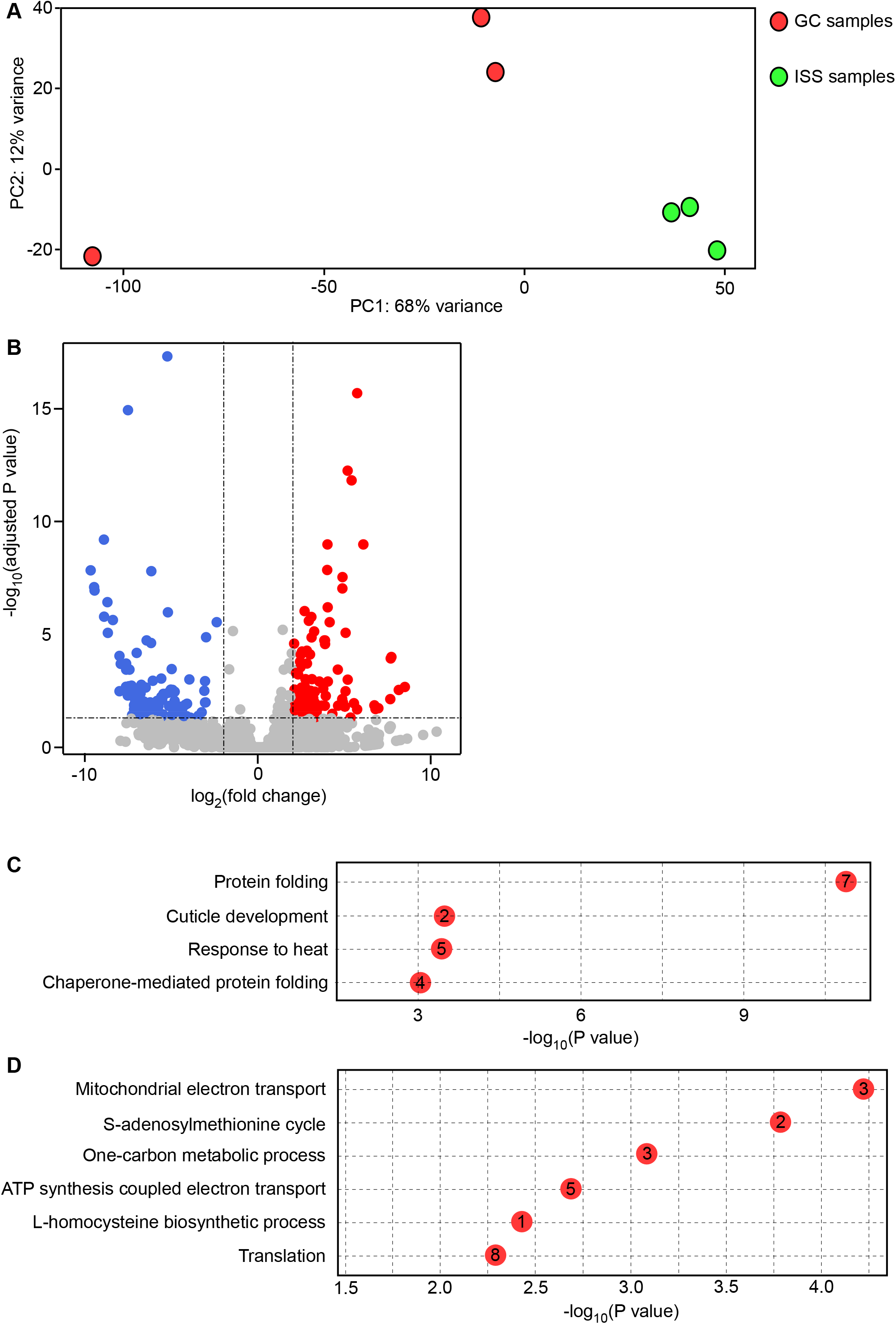
Transcriptomic alterations in active tardigrades exposed to conditions aboard the ISS. (A) Principal component analysis (PCA) of gene expression profiles from ISS-exposed and ground-control tardigrades. The plot demonstrates distinct separation between the two groups based on their transcriptomic profiles. Each group comprises three independent biological replicates, with each replicate consisting of approximately five hundred tardigrades per tube. (B) Volcano plot showing differentially expressed genes in ISS-exposed tardigrades relative to ground controls. Genes were considered significantly differentially expressed at an adjusted *P* < 0.05 and an absolute log₂ fold change ≥ 1. (C and D) Gene Ontology (GO) enrichment analysis of genes significantly upregulated (C) or downregulated (D) in ISS-exposed tardigrades relative to ground controls. Numbers within the circles indicate the number of differentially expressed genes associated with each GO term.

We next performed differential gene expression analysis to identify transcriptional changes in tardigrades exposed to the ISS environment relative to ground controls. This analysis identified 86 significantly upregulated genes (log₂FC ≥ 1) and 139 significantly downregulated genes (log₂FC ≤ −1) in ISS-exposed tardigrades relative to ground controls (adjusted P < 0.05) (Fig. 3B, Table S4). To investigate the biological processes associated with these transcriptional changes, we performed Gene Ontology (GO) enrichment analysis separately for the upregulated and downregulated DEGs (Table S5). Genes associated with protein folding and cellular responses to heat were significantly enriched among the upregulated DEGs. In contrast, the downregulated DEGs were enriched for genes associated with ATP synthesis, mitochondrial electron transport, one-carbon metabolism, and protein synthesis (Fig. 3 C and D). Collectively, these findings suggest that exposure to the ISS environment is associated with increased expression of genes involved in proteostasis and stress responses, accompanied by reduced expression of genes involved in energy metabolism and protein synthesis.

## DISCUSSION

The ability of tardigrades to survive extreme physical conditions has long fascinated astrobiologists. Several species have previously been exposed to space to assess their survival and reproductive capacity [9, 10, 22]. For instance, *Richtersius coronifer* (Richters, 1903) [23] and *Milnesium tardigradum* (Doyère, 1840) [24] were sent to low Earth orbit as part of the European Space Agency’s FOTON-M3 mission. Both species survived exposure to the vacuum of space but not to its ultraviolet radiation [8]. Similarly, *Macrobiotus richtersi* (Murray, 1911) [25] was part of the LIFE-TARSE experiment on FOTON-M3, where desiccated specimens showed survival comparable to ground controls after 12 days in orbit [11]. Notably, these missions did not involve reactivating dormant tardigrades in space or aboard the ISS, nor did they involve analyzing gene expression. Our study, using tardigrades, shows that a dormant multicellular animal can be activated beyond Earth at the ISS, followed by successful recovery and reproduction after return. Our results contribute to the current understanding of multicellular dormant life under spaceflight conditions.

The reduced survival of tardigrades observed in the ISS group likely reflects the cumulative effects of multiple mission-associated stresses. These include launch and re-entry, which subject the samples to high vibration and G-forces, as well as exposure to the spaceflight environment. In addition, rehydration under microgravity may have introduced variability because fluid behavior in microgravity conditions is governed predominantly by surface tension and capillary forces [26, 27]. This can potentially lead to non-uniform water distribution across the filter paper and variable hydration of dormant specimens. We acknowledge that a dedicated launch-stress control was not feasible under current mission constraints. Therefore, the current experimental design does not permit attributing our observations specifically to microgravity or other spaceflight-associated variables. Instead, the observed difference is likely due to the cumulative stress associated with transport to and from the ISS (launch, transport, residence aboard the ISS).

While vertebrates are far more complex than tardigrades and lack the capacity for dormancy, several mammals naturally enter hibernation, a state of prolonged torpor characterized by reduced metabolism and body temperature. Examples include the small Madagascar hedgehog (*Echinops telfairi*), the little brown bat (*Myotis lucifugus*), the thirteen-lined ground squirrel (*Ictidomys tridecemlineatus*), and the fat-tailed dwarf lemur (*Cheirogaleus medius*). Hibernation found in these animals is fundamentally different from the dormancy found in tardigrades and other less complex animals. However, both states are associated with increased tolerance to certain physiological stresses, although they arise through fundamentally different biological mechanisms [28, 29]. For example, certain hibernating mammals don’t exhibit musculoskeletal deterioration, which is a common problem faced by astronauts in space [30].

Gene expression analyses have revealed that the molecular programs controlling hibernation in these mammals are also present in humans, where they regulate key metabolic processes [31]. These studies suggest that some molecular programs associated with metabolic suppression are conserved in mammals [32], raising the possibility of artificially inducing a reversible, hibernation-like state termed synthetic torpor. Indeed, experimental approaches to induce such a state are underway. For example, experimental induction of torpor-like states in rodents has been explored using several approaches, including exposure to hydrogen sulfide (H₂S) [33], microinjection of GABA_A_ agonist muscimol into certain central nervous pathways [34], activation of the adenosine A1 receptors and activation of the hypothalamic Q neurons [35, 36]. Such studies have stimulated broader interest in reversible metabolic suppression for long-duration spaceflight missions [3–5, 28, 37–39].

Our transcriptomic analyses provided insights into the molecular adaptations of tardigrades to conditions aboard the ISS. Notably, the upregulation of genes involved in protein folding and chaperone-mediated processes suggests an enhanced capacity to maintain protein homeostasis and mitigate the accumulation of misfolded proteins, a common consequence of various environmental stresses [40]. In contrast, the downregulation of genes associated with multiple metabolic pathways suggests a shift in cellular priorities, potentially redirecting resources from energy-intensive metabolic processes toward cellular maintenance and damage repair. Such a response may represent an adaptive strategy to conserve energy and promote survival under stressful conditions, particularly when nutrient availability is limited. Importantly, the functions of several genes differentially expressed under ISS conditions remain unknown (Table S4). These genes represent promising candidates for future investigations aimed at identifying previously uncharacterized molecular mechanisms underlying stress tolerance and adaptation to extreme environments.

Similar to our study, previous studies involving ISS-exposed *Arabidopsis*, *Drosophila*, and bdelloid rotifers have reported increased expression/abundance of molecular chaperones and heat-shock proteins [41–43]. Together, these observations suggest that enhanced protein homeostasis represents a common cellular response to spaceflight-associated stresses across evolutionarily diverse organisms.

Further studies are required to understand the molecular and physiological mechanisms that enable *P. karunaadensis* to endure repeated cycles of dormancy and reactivation under conditions associated with spaceflight. Identifying the protective pathways and biomolecules involved could lead to strategies to preserve cellular integrity during prolonged metabolic arrest in other model organisms. Such insights may broaden understanding of biological tolerance to prolonged metabolic suppression under extreme conditions, advancing our understanding of life’s limits.

## METHODS

### Tardigrade culture and dormancy induction

The tardigrades used in this study were originally isolated from a moss sample collected from a concrete wall in Bengaluru, India. Detailed morphological features and phylogenetic analysis have been previously published [18]. Cultures were maintained in 35 mm Petri dishes coated with 2% agarose (Lonza, Switzerland) using KCM solution (7 mg KCl, 8 mg CaCl₂, and 8 mg MgSO₄ · 7H₂O per liter of water) as the medium. Cultures were kept in the dark at 20°C and fed *Caenorhabditis elegans* weekly. Axioscope upright light microscope (Carl Zeiss) with N-Achroplan 10x/0.25 Ph1 M27 objective was used for imaging.

To prepare for anhydrobiosis induction in the lab, tardigrades were starved for 24 h at 20°C. The animals were then transferred onto a Whatman grade-3 filter paper with 20 µL of reverse osmosis (RO) water. Anhydrobiosis was induced by first desiccating the animals at 20°C for 24 h, followed by a 48-h incubation at 25°C in a desiccator containing NaOH pellets (relative humidity ∼ 55%). This protocol facilitated the induction of dormancy (the anhydrobiotic tun state). For rehydration, 50 µL aliquots of RO water was added to the edges of the filter paper every 15 minutes over a 1-hour period (Fig. 2B). The filter paper was then flooded with RO water to dislodge the revived animals, which were subsequently returned to standard culture conditions described above.

### Genome sequencing

Approximately 2000 tardigrades were used for genomic DNA extraction with the QIAamp DNA Mini Kit (#51306, QIAGEN). The animals were subjected to multiple freeze-thaw cycles using dry ice and room-temperature incubation to facilitate disruption. Sterile ceramic beads and tissue lysis buffer were then added, and the samples were homogenized by bead beating for 2–3 min at 25 Hz using a TissueLyser. Proteinase K was subsequently added, and the samples were incubated for 2 h with shaking to ensure complete lysis. The remaining DNA extraction steps were performed according to the manufacturer’s instructions. Genomic DNA was eluted in nuclease-free water and quantified using a NanoDrop One spectrophotometer and a Qubit fluorometer. DNA integrity and fragment-size distribution were further assessed using an Agilent 4200 TapeStation with Genomic DNA ScreenTape.

The genome was sequenced using a hybrid approach integrating Illumina short reads (NextSeq 2000) and Oxford Nanopore long reads (PromethION, Flowcell R10.4.1). Reads from both platforms were filtered with Kraken2 against the GTDB 2024 database to remove bacterial contamination. Flye assembler was used to generate the initial assembly from long reads, followed by refinement with short reads using Pilon. Due to the high volume of Illumina reads, which could potentially impact assembly polishing efficiency, the reads were downsampled using BBnorm before the polishing stage. The assembly statistic was generated by QUAST. Assembly completeness was assessed using BUSCO v6.0.0 in genome mode, which identifies the presence of a predefined set of universal single-copy orthologs (BUSCOs). The metazoa_odb12 lineage dataset was used for this evaluation. We used OmicsBox to perform the full annotation workflow, including masking repetitive elements, predicting coding genes, and assigning functional annotations.

### Exposure to ultra-low temperature and pressure

The dormant tardigrades were cooled to ∼12 mK using a cryogen-free dilution refrigerator (Bluefors LD250). They were placed inside a small chamber made from oxygen-free high-conductivity copper and securely mounted to the mixing-chamber plate. The dilution refrigerator provides not only an ultracold thermal environment but also an ultralow vacuum (∼10⁻^12^ bar) at these cryogenic temperatures.

### Exposure to radiation

Tardigrades were exposed to gamma radiation sourced from the cobalt-60 isotope. The dose rate was 0.82 to 0.87 Gy/sec. Exposure time was determined by the required dosage (1000 to 6000 Gy). Control samples (0 Gy) were taken to the facility but were not exposed to radiation.

### A Science Verification Test (SVT)

SVT was conducted prior to the mission to validate the experimental setup, sample preparation, activation procedure, imaging protocol, and overall operational sequence. The SVT also ensured that the procedure could be reliably executed within the available operational constraints while yielding the intended biological observations. Here, tardigrades were maintained under laboratory conditions, using the same time intervals as for ISS and Ground control samples. This group served as an additional control (Laboratory control).

### Experiments at ISS

The experiment was initiated with three biological replicates per group, each consisting of approximately 90 tardigrades in a 35-mm dish. This number was selected to ensure sufficient animals for subsequent survival and reproductive analyses while remaining compatible with the available filter area and the operational constraints of the spaceflight experiment. One ISS biological replicate could not be analyzed due to damage, leaving two evaluable ISS replicates comprising approximately 180 tardigrades.

On board the ISS, the experiment was conducted at the Maintenance Work Area (MWA) in Node 2, where the temperature was maintained at approximately 22°C. The tardigrades were not provided with additional radiation shielding and were exposed to a radiation dose of 0.1863 to 0.2527 mGy d⁻¹. On day 6^th^ aboard ISS, dormant tardigrades were reactivated by the astronaut through gentle addition of 50 μL RO water, as described above (Fig. 2B). One hour after the final addition of water, the samples were examined using a Keyence BZX800 microscope equipped with a Plan Fluorite 4× PH objective (NA 0.13; working distance, 16.50 mm). Bright-field monochrome videos were recorded at 40 frames s⁻¹ over a total observation period of 15 min. On the same day, RNAlater, an aqueous RNA-stabilizing reagent, was added to active tardigrades and frozen at -80°C for transcriptome analysis.

Ground control tardigrades experienced the same environmental conditions as ISS tardigrades, except for spaceflight and ISS-associated microgravity and radiation. They were even taken from our lab to the Kennedy Space Center along with ISS samples and brought back. The storage duration was identical. The rehydration was done exactly at the same time as the ISS samples. We were in real-time communication with ISS astronauts on that day to ensure rehydration and RNAlater addition were performed at the same time and in the same manner.

Each Petri dish was treated as one independent biological replicate. Each dish contained ∼90 dormant tardigrades. Recovery, survival, egg laying, and offspring production were quantified at the dish level. One of the Petri dishes was damaged during transport from ISS and was excluded from the analysis. Statistical comparisons between groups were performed using unpaired two-tailed t-tests with Welch’s correction. Because the number of biological replicates was limited by mission logistics, statistical interpretation was performed cautiously, and non-significant differences were interpreted descriptively rather than as evidence of equivalence. Data are presented as mean ± SE, with each point in the graphs representing one biological replicate (dish).

### RNA isolation, sequencing, and quality control

The tardigrade samples were washed with nuclease-free water and lysed in 600 µL of TRIzol reagent using repeated freeze-thaw cycles. To ensure thorough tissue disruption and homogenization, the samples were subjected to bead beating using a TissueLyser at 20 Hz for 1 min. The resulting homogenates were centrifuged to remove insoluble debris, and the clarified supernatants were collected for RNA isolation using Qiagen miRNeasy Mini kit (# 1038703).

The purified RNA was subjected to next-generation sequencing on an Illumina platform. The quality of the raw sequencing reads was assessed using FastQC. Adapter sequences and low-quality bases (Phred quality score < 30) were removed using fastp, and reads shorter than 50 nucleotides after trimming were discarded. To minimize microbial contamination, the quality-filtered reads were screened using Kraken2 against a reference database derived from the Genome Taxonomy Database (GTDB, 2024 release), and reads classified as bacterial or archaeal were excluded. The remaining reads were processed using SortMeRNA (v4.3.6) against reference ribosomal RNA (rRNA) databases to identify and remove rRNA-derived sequences. The RNA sequencing data have been deposited in the NCBI Sequence Read Archive (SRA) under BioProject accession number PRJNA1540695.

### De novo transcriptome assembly and functional annotation

The resulting high-quality, non-rRNA paired-end reads were used for de novo transcriptome assembly using Trinity (v2.x), with the parameters --seqType fq, --max_memory 50G, and --CPU 16. The assembly yielded 84,077 putative genes, identified using Trinity’s gene-level nomenclature (TRINITY_DN*_c*_g*). Potential protein-coding regions within the assembled transcripts were predicted using TransDecoder. Functional annotation was performed using the Trinotate pipeline, incorporating BLAST-based sequence similarity searches against the UniProtKB/Swiss-Prot protein database, HMMER-based identification of conserved protein domains against the Pfam database, and SignalP-based prediction of signal peptides.

### Differential Expression Analysis

Differential gene expression analysis between ISS-exposed and ground-control samples was performed using DESeq2 (v1.42). Raw gene-level read counts were modeled using a negative binomial generalized linear model, with the experimental condition (ISS versus ground control) specified as the explanatory variable (`design = ∼ condition`). Differences in gene expression between the two conditions were assessed using Wald tests. Library size differences were accounted for by estimating size factors using DESeq2’s median-of-ratios normalization method. The resulting *P*-values were adjusted for multiple testing using the Benjamini–Hochberg procedure to control the false discovery rate (FDR). Genes with an adjusted *P*-value < 0.05 and an absolute log₂ fold change ≥ 1 were considered differentially expressed.

### Principal Component Analysis (PCA)

This was performed using variance-stabilizing transformed (VST) gene expression counts. The first two principal components (PC1 and PC2) explained 68% and 12% of the total variance, respectively. PC1 clearly separated the ground-control (GC) and ISS-exposed samples, indicating that the experimental condition was associated with the major source of variation in gene expression. Although GC replicates appeared more dispersed than ISS replicates along PC2, this axis accounted for a relatively smaller proportion of the total variance. To determine whether the two groups differed in overall multivariate dispersion, we performed a permutation-based test for homogeneity of multivariate dispersions (PERMDISP). The analysis did not detect a statistically significant difference in within-group dispersion (F(1,4) = 1.36, permutation *P* = 0.40). However, given the limited number of biological replicates (n = 3 per group), the statistical power to detect differences in dispersion was low. Overall, PCA revealed a clear separation between GC and ISS samples along the principal axis of variation.

### Gene Ontology (GO) Enrichment Analysis

This analysis was performed using topGO (v2.54) to identify biological functions overrepresented among differentially expressed genes. Upregulated and downregulated genes were analyzed separately across the three GO categories: Biological Process (BP), Molecular Function (MF), and Cellular Component (CC). Overrepresentation of GO terms was assessed using Fisher’s exact test with the ‘weight01’ algorithm. For each enriched GO term, the contributing differentially expressed genes were identified by intersecting the genes associated with that term, retrieved using ‘genesInTerm()’, with the significant gene set obtained using ‘sigGenes()’ from the corresponding topGO object. GO terms with a nominal *P*-value < 0.01 were retained for further analysis.

## Acknowledgements

This study was conducted as part of the experiment titled “Survival, revival, reproduction and transcriptome of the Eutardigrade *Paramacrobiotus* Sp. BLR strain under microgravity”, selected by ISRO’s Human Space Flight Centre (HSFC) for ISRO-NASA Joint Mission to the ISS (Axiom-4 mission). The authors are grateful to HSFC/ISRO for enabling this study during the Axiom-4 mission. We thank Capt. Shubhanshu Shukla, Anurag K. Sinha, and Tushar Phadnis for their valuable technical assistance. This work was supported by the Indian Space Research Organization’s Space Technology Cell (ISRO-STC) at the Indian Institute of Science, Bengaluru (ISTC/BES/SME/480 and ISTC/BES/SME/513). Funds from Swarnajayanti Fellowship (DST/SJF/LSA-04/2019-20 to SME) given by the Department of Science and Technology (DST), EMBO Global Investigator Network, and DST Funds for Improvement of S&T infrastructure are acknowledged.

## SUPPLEMENTARY INFORMATION

**Legend to Video S1.** Activation of dormant tardigrades at ISS. The video shows the movement of tardigrades after activation. Samples were imaged one hour after the activation using a Keyence BZ-X800 microscope. A Plan Fluorite 4× PH objective (NA 0.13; working distance 16.50 mm) was used. Brightfield monochrome videos were acquired at 40 frames per second.

**Table S1.** Genome assembly statistics.

| **Feature** | **Values** |
|---|---|
| Number of scaffolds | 4100 |
| Number of contigs | 4100 |
| Largest contig | 4722118 |
| Total length | 127879965 |
| GC (%) | 47.11 |
| N50 | 103kbp |
| Minimum Length | 447 |
| Maximum Length | 1602447 |

**Table S2.**
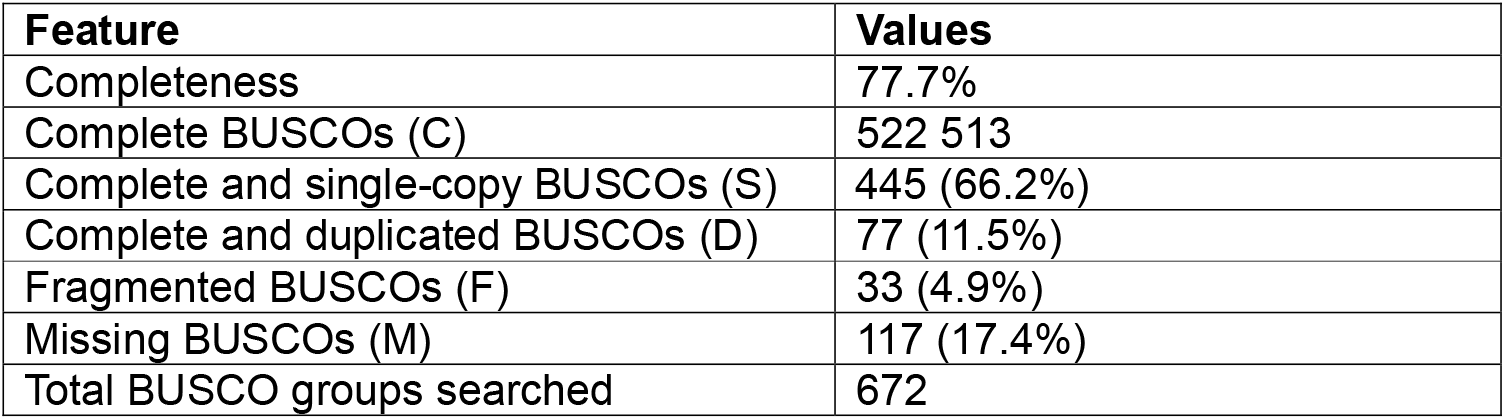
The summary of assembly evaluation by BUSCO.

| **Feature** | **Values** |
|---|---|
| Completeness | 77.7% |
| Complete BUSCOs (C) | 522 513 |
| Complete and single-copy BUSCOs (S) | 445 (66.2%) |
| Complete and duplicated BUSCOs (D) | 77 (11.5%) |
| Fragmented BUSCOs (F) | 33 (4.9%) |
| Missing BUSCOs (M) | 117 (17.4%) |
| Total BUSCO groups searched | 672 |

**Table S3.** Summary of BLASTn results^#^ for selected predicted marker genes.

| Marker gene from the new strain | Best match | Query Cover | E-value | Identity |
|---|---|---|---|---|
| 60S ribosomal protein L18a-like | *Paramacrobiotus metropolitanus* 60S ribosomal protein L18a-like (LOC129597494) | 100% | 0 | 89.70% |
| Histone H2A | *Paramacrobiotus metropolitanus* histone H2A (LOC129582550) | 100% | 4e-150 | 88.30% |
| cytochrome c oxidase subunit 5A, mitochondrial-like | *Paramacrobiotus metropolitanus* cytochrome c oxidase subunit 5A, mitochondrial-like (LOC129584881) | 100% | 9e-126 | 87.19% |
| cytochrome c oxidase subunit 4 isoform 1, mitochondrial-like | *Paramacrobiotus metropolitanus* cytochrome c oxidase subunit 4 isoform 1, mitochondrial-like (LOC129595091) | 100% | 1e-160 | 83.63% |
| tubulin beta chain- like | *Paramacrobiotus metropolitanus* tubulin beta chain-like (LOC129586219) | 49% | 0 | 86.83% |
| homeobox protein SIX3-like | *Paramacrobiotus metropolitanus* homeobox protein SIX3-like (LOC129581640) | 86% | 0 | 88.63% |
| RNA polymerase II subunit RPB1 | *Paramacrobiotus metropolitanus* RNA polymerase II subunit RPB1 (LOC129586423) | 100% | 0 | 84.82% |
| 18S_rRNA | *Macrobiotus richtersi* group 2 18S ribosomal RNA gene, partial sequence<br>*Paramacrobiotus metropolitanus* TYO gene for 18S ribosomal RNA, partial sequence | 100%<br>100% | 0<br>0 | 99.37%<br>98.97% |
| 28S_rRNA | *Paramacrobiotus metropolitanus* large subunit ribosomal RNA (LOC129600597). | 100% |  | 99.39% |

<sup>#</sup> The top-most match for every gene in the BLASTn analysis is shown

Table S4: List of differentially expressed genes (uploaded separately as an Excel file)

Table S5: Results of Gene Ontology (GO) analysis (uploaded separately as an Excel file)

